# ASSESSING TARGET SPECIFICITY OF THE SMALL MOLECULE INHIBITOR MARIMASTAT TO SNAKE VENOM TOXINS: A NOVEL APPLICATION OF THERMAL PROTEOME PROFILING

**DOI:** 10.1101/2023.10.25.564059

**Authors:** Cara F. Smith, Cassandra M. Modahl, David Ceja-Galindo, Keira Y. Larson, Sean P. Maroney, Lilyrose Bahrabadi, Nicklaus P. Brandehoff, Blair W. Perry, Maxwell C. McCabe, Daniel Petras, Bruno Lomonte, Juan J. Calvete, Todd A. Castoe, Stephen P. Mackessy, Kirk C. Hansen, Anthony J. Saviola

## Abstract

New treatments that circumvent the pitfalls of traditional antivenom therapies are critical to address the problem of snakebite globally. Numerous snake venom toxin inhibitors have shown promising cross-species neutralization of medically significant venom toxins *in vivo* and *in vitro*. The development of high-throughput approaches for the screening of such inhibitors could accelerate their identification, testing, and implementation, and thus holds exciting potential for improving the treatments and outcomes of snakebite envenomation worldwide. Energetics-based proteomic approaches, including Thermal Proteome Profiling (TPP) and Proteome Integral Solubility Alteration (PISA), assays represent “deep proteomics” methods for high throughput, proteome-wide identification of drug targets and ligands. In the following study, we apply TPP and PISA methods to characterize the interactions between venom toxin proteoforms in *Crotalus atrox* (Western Diamondback Rattlesnake) and the snake venom metalloprotease (SVMP) inhibitor marimastat. We investigate its venom proteome-wide effects and characterize its interactions with specific SVMP proteoforms, as well as its potential targeting of non-SVMP venom toxin families. We also compare the performance of PISA thermal window and soluble supernatant with insoluble precipitate using two inhibitor concentrations, providing the first demonstration of the utility of a sensitive high-throughput PISA-based approach to assess the direct targets of small molecule inhibitors for snake venom.

## Introduction

Snakebite is a global public health problem that disproportionately affects impoverished communities in rural tropical and subtropical regions. Annual estimates suggest that snakebite affects 1.8–2.7 million people worldwide, causing >138,000 deaths and leaving an even larger number of victims suffering permanent disabilities (1), which has led to the designation of snakebite as a neglected tropical disease by the World Health Organization (WHO; 2,3). Snake venoms are complex toxic cocktails of proteins and peptides derived from more than a dozen gene families, many of which have undergone duplication to generate multiple functionally diverse paralogs and associated proteoforms in the venom of a single species (4–6). While substantial variation exists in the relative mass and functional activity of venom proteins and peptides, most of these toxins have evolved to target and disrupt numerous bodily systems (7–10). Adding to the complexity of snake venoms, the most medically relevant toxin families tend to be the most diverse with many paralogs and associated proteoforms displaying moderate-to-high sequence similarity but in many cases exhibiting a spectrum of distinct biological effects (7,9,11–13). One of these families, snake venom metalloproteases (SVMPs), is ubiquitous across snake species (but is particularly abundant in viperid venoms) and is responsible for many of the life-threatening pathologies that result from snake envenomation, including local and systemic hemorrhage, and tissue destruction (14–18).

The substantial morbidity and mortality resulting globally from snakebite may seem surprising considering that antivenoms (whole IgG molecules, Fab or F(ab′)_2_ fragments from venom-immunized animals) are often highly effective at recognizing and neutralizing the major toxic components of a venom (19,20). A major challenge to antivenom efficacy, however, is the significant variation in venom composition that occurs at phylogenetic (21–26), ontogenetic (27–30), and geographic or population scales (31–36). As a consequence, antivenoms are most effective against the snake species whose venom was utilized during production and are often inadequate at recognizing venom components of different or even closely related snake species (25). Further, geographic venom variation may even result in poor neutralization within the same species when snakes used in antivenom production were sourced from a different location (33). Antivenoms also tend to be more effective at neutralizing systemic effects, but less effective at neutralizing anatomically localized manifestations of envenomation, which can result in permanent tissue damage and disfigurement (1,37–41). The storage, accessibility, and administration of antivenom also pose significant practical challenges in rural areas where it is needed most (10,37,42–48). These hurdles are further compounded by the excessive effort and cost of producing antivenoms for any one geographically-relevant set of venomous snake species.

While the use of polyvalent antivenoms has been the mainstay treatment for snake envenomation, the development of non-immunological treatments that circumvent the limitations of antivenoms has been prioritized as a goal to address the impacts of snake envenomation globally by the WHO (3). Recent applications of small molecule inhibitors against medically significant toxins have yielded promising preclinical results and these inhibitors have broad potential as supplemental therapies in combination with standard treatments (37,49–54). These inhibitors have a number of advantages over current antivenom therapies including better peripheral tissue distribution, higher shelf stability, a higher safety profile, the ability for pre-hospital oral or topical administration, and greater affordability (55). The use of novel high-throughput approaches for the testing of venom toxin inhibitors and the identification of their targets could accelerate the implementation of effective small molecule inhibitors, with the long-term potential of improving the treatment and outcome of snakebite envenomation globally.

Numerous studies have examined the effects of various small molecule inhibitors on the biological activities of venoms *in vitro* and the neutralization capacity of these inhibitors *in vivo* (49–51,53,56–59). A repurposed low-specificity matrix metalloprotease inhibitor, marimastat, has shown effective neutralization of SVMP-rich venoms across multiple venomous snake species by preventing both local and systemic toxicity (53), decreasing hemotoxic venom effects (50,51,53,60,61), reducing SVMP-induced cytotoxicity (56,60), and inhibiting extracellular matrix degradation (62). Because marimastat has previously progressed to clinical trials as a cancer treatment, its safety profile has already been determined, accelerating its development as a potential snakebite treatment (63,64). When administered with other small molecule inhibitors, marimastat has shown *in vivo* neutralization of lethal toxicity and dermonecrosis in murine models (50,56). Each of these studies assesses the downstream effects of inhibitor action *in vivo* or *in vitro* by measuring changes to biological activity or survival; however, to our knowledge no studies exist using a direct assessment of venom-wide target-ligand interactions between venom toxins and small molecule inhibitors.

Thermal Proteome Profiling (TPP) is a prominent energetics-based proteomic approach for identifying the molecular targets of drugs. TPP builds upon the concept that a protein’s physicochemical properties are altered through interactions with extrinsic factors (e.g., other proteins, therapeutic drugs, metabolites) making it more or less resistant to thermal-induced denaturation (65,66). Traditional TPP assays were centered on the principle that unbound proteins tend to denature and become insoluble when subjected to increasing temperatures, whereas proteins stabilized through interactions with extrinsic factors often exhibit increased thermal stability and remain in solution (65,67–69). Identifying and quantifying such solubility changes via mass spectrometry can be used to infer direct or indirect interactions between a given compound and its protein targets (70).

Recently, the Proteome Integral Solubility Alteration (PISA) assay has emerged as a powerful strategy that retains the breadth and sensitivity of TPP but with a significant reduction in sample preparation and analysis time (70,71). PISA represents a “deep proteomics” method for high throughput proteome-wide target identification of ligands, with improved target discovery and higher statistical significance for target candidates (70–72). In a PISA assay, samples are subjected to heat across a temperature gradient (as in TPP) but are subsequently pooled prior to analysis (70,71). Rather than generating melt curves to determine exact melting temperatures, PISA compares overall abundance of each measured peptide between controls and treatments to detect differences in melting properties when a compound of interest is added. This methodology allows multiple variables to be altered simultaneously (e.g., concentration, temperature) in a high-throughput manner. It has recently been shown that heat-treating within a smaller temperature window can improve sensitivity and target discovery with PISA (73). TPP, PISA, and related methods derived from the same principles have been used to discover drug targets, antibiotic targets, and mechanisms of antibiotic resistance (65,66,69,74). In our specific context, these proteomic techniques applied to the development of envenomation treatments hold strong potential to provide rapid and high-throughput characterization of small molecule venom toxin inhibitors by determining their direct targets across diverse venom toxin protein families, accelerating identification of novel inhibitors.

Here, we apply TPP and PISA methods to characterize the physical interactions between the SVMP inhibitor marimastat and toxin proteoforms of *Crotalus atrox* (Western Diamondback Rattlesnake) venom. First, we determined toxin proteoform presence and abundance in the venom of this well-studied species and used TPP to characterize the venom meltome by determining venom protein family-level and specific proteoform-level thermal characteristics. Next, we performed PISA experiments within two different thermal windows to assess protein thermal stability changes upon inhibitor addition to identify specific proteoform targets of the small molecule inhibitor marimastat. Because of the previously characterized differences in signal-to-noise ratio between supernatant and pellet in PISA experiments (75,76), we investigate and compare the targets identified in both the soluble and insoluble fractions. Our results demonstrate that a PISA-based approach can provide rapid, highly sensitive, and robust inferences for the unbiased proteome-wide screening of venom and inhibitor interactions.

## Methods

### Venom and inhibitors

*Crotalus atrox* (Western Diamondback rattlesnake) venom was obtained by manual extraction from snakes housed at the University of Northern Colorado (UNC) Animal Facility (Greeley, CO), in accordance with UNC-IACUC protocols. Venoms were lyophilized and stored at -20°C until use. Venoms were reconstituted at a concentration of 2 mg/mL and protein concentration was determined on a Nanodrop™ using the Absorbance 280 program. The small molecule matrix metalloprotease inhibitor marimastat ((2*S*,3*R*)-*N*4-[(1*S*)-2,2-Dimethyl-1- [(methylamino)carbonyl]propyl]-*N*1,2-dihydroxy-3-(2-methylpropyl)butanediamide, >98%, Cat no.: 2631, Tocris Bioscience) was reconstituted in ddH_2_0 at a concentration of 1.5 mM and stored at −20°C.

### Venom gland transcriptomics

An adult *C. atrox* was collected in Portal, AZ under collecting permit 0456, and maintained in the UNC Animal Facility. Four days following manual venom extraction, the *C. atrox* was humanely euthanized and venom glands removed (IACUC protocol no. 9204). Approximately 70 mg of tissue, originating from both left and right venom glands, was homologized. Total RNA was isolated from homologized venom gland tissue using the previously described TRIzol (Life Technologies, C.A. U.S.A.) protocol for venom glands (77,78) A NEBNext Poly(A) mRNA Magnetic Isolation Module (New England Biolabs, MA, U.S.A) was used to select mRNA from 1 µg of total RNA, and the NEBNext Ultra RNA library prep kit (New England Biolabs, MA, U.S.A) manufacture’s protocol followed to prepare the sample for Illumina® RNA-sequencing (RNA-seq). During library preparation, products within the 200-400 bp size range were selected by solid phase reversible immobilization with the Agencourt AMPure XP reagent (Beckman Coulter, C.A., U.S.A.) and PCR amplification consisted of 12 cycles. Final quantification of the RNA-seq library was done with the Library Quantification Kit for Illumina® platforms (KAPA Biosystems, M.A, U.S.A.). The *C. atrox* venom gland RNA-seq library was then checked for proper fragment size selection and quality on an Agilent 2100 Bioanalyzer, equally pooled with eight other unique barcoded RNA-seq libraries and sequenced on 1/8^th^ of an Illumina® HiSeq 2000 platform lane at the UC Denver Genomics core to obtain 125 bp paired-end reads.

To produce a comprehensive venom gland transcriptome database for *C. atrox,* two RNA-seq libraires were *de novo* assembled, the first from the *C. atrox* RNA-seq library detailed above and the second from a Texas locality *C. atrox* with reads available on the National Center for Biotechnology Information server (SRR3478367). Low quality reads were trimmed and adaptors removed using Trimmomatic (79) with a sliding window of 4 nucleotides and a threshold of phred 30. Reads were then assessed with FastQC (Babraham Institute Bioinformatics, U.K.) to confirm that all adapters and low-quality reads were removed before *de novo* assembly. Three *de novo* assemblers were used in combination to produce a final, high-quality assembly: i). first, a Trinity (release v2014-07-17) genome-guided assembly was completed using default parameters and Bowtie2 (v2.2.6) (80) aligned reads to the *C. atrox* genome (provided by Noah Dowell (81)), ii) a second *de novo* assembly was completed with the program Extender (k-mer size 100) (82), performed with the same parameters as used for other snake venom glands (83), and with merged paired-end reads, merged with PEAR (Paired-End read mergeR v0.9.6; default parameters) (84), as seed and extension inputs, iii) a third *de novo* assembly was completed with VT Builder using default settings (85). From a concatenated fasta file of all three assemblies, coding contigs were then identified with EvidentialGene (downloaded May 2018) (86) and redundant coding contigs and those less than 150 bps were removed with CD-HIT (87,88). Reads were aligned with Bowtie2 to coding contigs and abundances determined with RSEM (RNA-seq by Expectation-Maximization; v1.2.23) (89). Contigs less than 1 TPM (Transcript Per Million) were filtered out, and the remaining contigs annotated with Diamond (90) BLASTx (E-value 10^-05^ cut-off) searches against the NCBI non-redundant protein database. Transcripts were identified as venom proteins after each was manually examined to determine if the resulting protein was full-length, shared sequence identity to a currently known venom protein, and contained a shared signal peptide sequence with other venom proteins within that superfamily. This transcript set was also filtered through ToxCodAn (91) as a final toxin annotation check, and the resulting translated toxins used as a custom database for mass spectrometry.

### Venom Meltome Generation

Thermal profiling assays were carried out following previously described methods (74,92). Venom (1 μg/μL) was assayed in duplicate and divided into 10 aliquots of 20 μL and transferred to 0.2 mL PCR tubes. Each aliquot was individually heated at a fixed temperature over the range of 37° to 75°C for 3 min in a Bioer LifeECO™ thermal cycler (Figure 1a). Samples were allowed to aggregate at room temperature for one minute and then placed on ice. Precipitated proteins were removed by centrifugation at 21,000 x g for 45 min and the supernatant was removed and subjected to sodium dodecyl sulfate polyacrylamide gel electrophoresis (SDS-PAGE) and sample preparation for mass spectrometric analysis.

**Figure 1.**
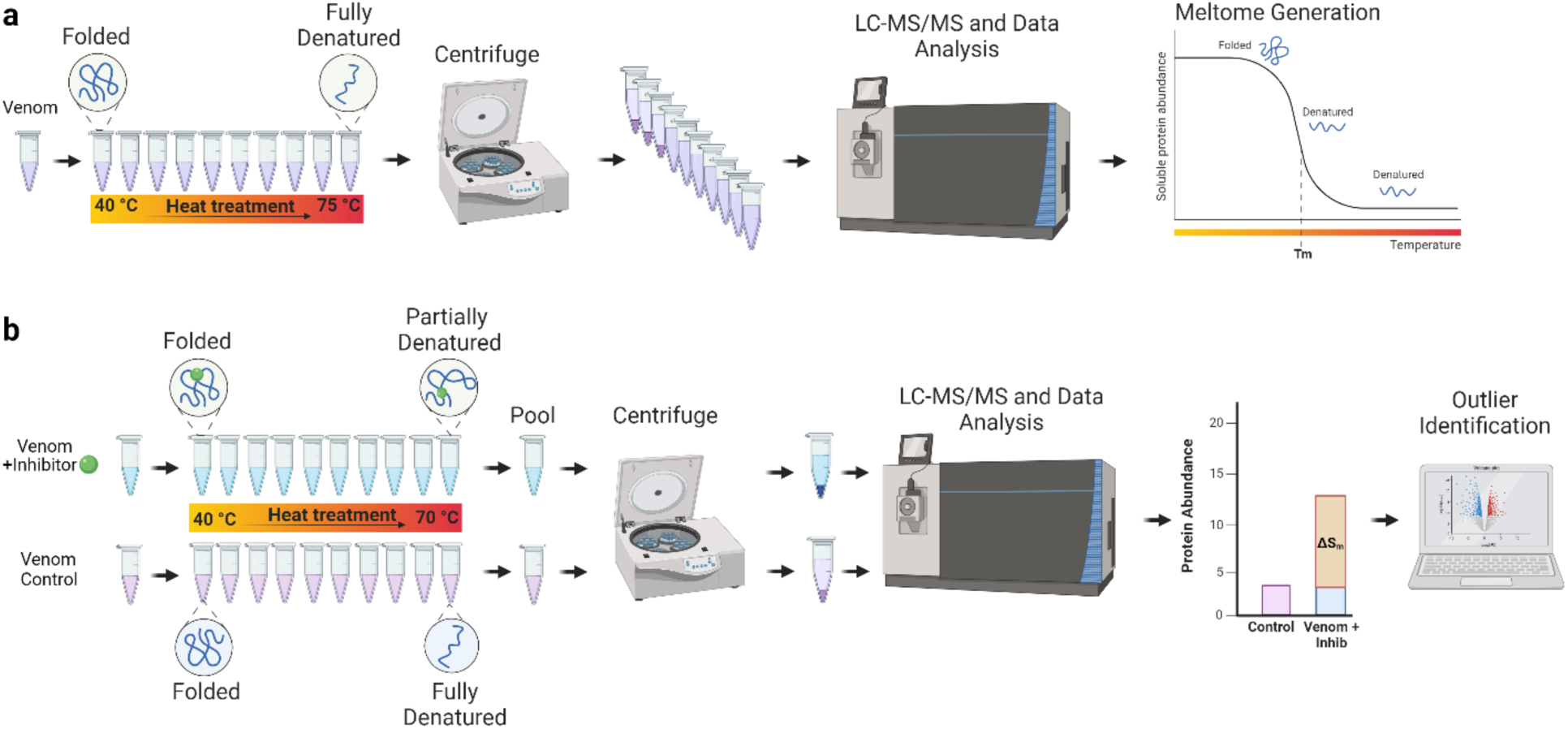
TPP (a) and PISA (b) workflows. a) In TPP experiments, samples are heated between 40 – 70°C and centrifuged to pellet denatured proteins. Samples are reduced, alkylated, and trypsin digested and analyzed with LC-MS/MS for protein identification. Melting curves are generated in ProSAP using unique intensity for each protein identified. *T_m_* = melting temperature of 50% of population. b) In PISA, venom is incubated for 30 minutes at 37°C with an inhibitor or alone. Samples are heated from 40-70°C, and pooled before centrifuging to pellet insoluble material. Samples are prepared as mentioned above and analyzed via LC-MS/MS for protein identification. To identify inhibitor targets, unique intensity is used to calculate SAR values for each protein followed by identification of significant outliers.

### Inhibitor PISA Assays

Venom (1 μg/μL) was incubated with two previously explored concentrations of marimastat, 15 μM or 150 μM (53,61), or a vehicle control (ddH_2_O) for 30 min at 37°C. Each sample was then divided into 12 aliquots of 20 μL in 0.2 mL PCR tubes. Each aliquot was individually heated at a different temperature from 40 to 70°C for 3 min in a Bioer LifeECO™ Thermal cycler (Figure 1b), allowed to cool at room temperature for one min, and placed on ice. An equal volume of sample from each temperature point was pooled and centrifuged at 21,000 x g for 45 min at 4°C to separate the soluble fraction from insoluble denatured proteins (70). Because of the previously characterized differences in performance between supernatant and pellet, we investigated both fractions (75,76). Approximately 30 μg of soluble protein (based on control samples) was collected and prepared for mass spectrometry and 20 μg was used for gel electrophoresis. PISA assays for each condition were performed in triplicate. Because selection of a narrower temperature window for heat denaturation has been shown to increase sensitivity of the PISA assay (73), we also performed a temperature gradient denaturation with 5 temperatures (selected based on SVMP family-level *T_m_* values) from 56 to 60°C. Samples were pooled and processed as described above.

### High-Performance Liquid Chromatography (HPLC)

One mg of venom incubated with either 150 μM marimastat or a vehicle control (ddH_2_O) was subjected to reverse phase HPLC after heat treatment using a Waters system, Empower software, and a Phenomenex Jupiter C_18_ (250 × 4.6 mm, 5 µm, 300 Å pore size) column as outlined in Smith and Mackessy (93). Proteins/peptides were detected at 280 nm and 220 nm with a Waters 2487 Dual λ Absorbance Detector. Fractions corresponding to each peak were then frozen at -80°C overnight, lyophilized, and then separated with SDS-PAGE as previously described (93). Percent peak area and peak height at 280 nm were recorded as a proxy for relative toxin abundance.

### Sodium dodecyl sulfate polyacrylamide gel electrophoresis (SDS-PAGE)

SDS-PAGE materials were obtained from Life Technologies, Inc. (Grand Island, NY, USA). Dithiothreitol (DTT)-reduced venom (20 µg) or lyophilized protein (approximately 5 µg – reverse-phase high-performance liquid chromatography (RP-HPLC) fractionated) was loaded into wells of a NuPAGE Novex Bis-Tris 12% acrylamide Mini Gel and electrophoresed in MES buffer under reducing conditions for 45 min at 175 V; 7 µL of Mark 12 standards were loaded for molecular weight estimates. Gels were stained overnight with gentle shaking in 0.1% Coomassie brilliant blue R-250 in 50% methanol and 20% acetic acid (v/v) and destained in 30% methanol, 7% glacial acetic acid (v/v) in water until background was sufficiently destained (approximately 2 hours). Gels were then placed in storage solution (7% acetic acid, v/v) for several hours with gentle shaking at room temperature and imaged on an HP Scanjet 4570c scanner.

### Sample preparation for Liquid chromatography-tandem mass spectrometry (LC-MS/MS)

The volume of supernatant corresponding to 30 μg of unmelted (soluble) protein was dried in a speed vacuum and redissolved in 8 M urea/0.1 M Tris (pH 8.5) and reduced with 5 mM TCEP (tris (2-carboxyethyl) phosphine) for 20 min at room temperature. Samples were then alkylated with 50 mM 2-chloroacetamide for 15 min in the dark at room temperature, diluted 4-fold with 100 mM Tris-HCl (pH 8.5), and trypsin digested at an enzyme/substrate ratio of 1:20 overnight at 37°C. To stop the reaction, samples were acidified with formic acid (FA), and digested peptides were purified with Pierce^TM^ C18 Spin Tips (Thermo Scientific #84850) according to the manufacturer’s protocol. Samples were dried in a speed vacuum and redissolved in 0.1% FA.

Electrophoretic protein bands subjected to LC-MS/MS were excised from Coomassie-stained gels, destained, and subjected to in-gel reduction, alkylation, and overnight trypsin digestion as previously described (94). Following the overnight digestion, samples were acidified with 5% formic acid (FA) and tryptic peptides were extracted in 30 µl of 50% acetonitrile /1% FA. Digests were dried in a vacuum centrifuge and redissolved in 0.1% FA for mass spectrometry.

### Nano liquid chromatography tandem mass spectrometry

Nano Liquid Chromatography tandem mass spectrometry (Nano-LC-MS/MS) was performed using an Easy nLC 1000 instrument coupled with a Q-Exactive™ HF Mass Spectrometer (both from ThermoFisher Scientific). Approximately 3 μg of digested peptides were loaded on a C_18_ column (100 μm inner diameter × 20 cm) packed in-house with 2.7 μm Cortecs C_18_ resin, and separated at a flow rate of 0.4 μL/min with solution A (0.1% FA) and solution B (0.1% FA in ACN) under the following conditions: isocratic at 4% B for 3 min, followed by 4%-32% B for 102 min, 32%-55% B for 5 min, 55%-95% B for 1 min and isocratic at 95% B for 9 min. Mass spectrometry was performed in data-dependent acquisition (DDA) mode. Full MS scans were obtained from *m/z* 300 to 1800 at a resolution of 60,000, an automatic gain control (AGC) target of 1 × 10^6^, and a maximum injection time (IT) of 50 ms. The top 15 most abundant ions with an intensity threshold of 9.1 × 10^3^ were selected for MS/MS acquisition at a 15,000 resolution, 1 × 10^5^ AGC, and a maximal IT of 110 ms. The isolation window was set to 2.0 *m/z* and ions were fragmented at a normalized collision energy of 30. Dynamic exclusion was set to 20 s.

### Analysis of mass spectrometry data

Fragmentation spectra were interpreted against a custom protein sequence database generated from the assembly of *C. atrox* venom gland transcriptome data (described above) that was combined with UniProt entries of all toxins found in the *C. atrox* venom proteome reported by Calvete et al. (95) using MSFragger within the FragPipe computational platform (96,97). Reverse decoys and contaminants were included in the search database. Cysteine carbamidomethylation was selected as a fixed modification, oxidation of methionine was selected as a variable modification, and precursor-ion mass tolerance and fragment-ion mass tolerance were set at 20 ppm and 0.4 Da, respectively. Fully tryptic peptides with a maximum of 2 missed tryptic cleavages were allowed and the protein-level false discovery rate (FDR) was set to < 1%. The relative abundance of major snake venom toxin families was compared across samples using sum-normalized total spectral intensity (98).

### Analysis of TPP data

Protein melting curves were generated by fitting sigmoidal curves to relative protein abundances using the Protein Stability Analysis Pod (ProSAP) package (99). The temperature at which relative protein abundance reached 50%, *T_m_* (melting temperature), was determined in ProSAP by normalizing intensity to the lowest temperature (37°C), followed by normalization to the most thermostable proteins as previously described (76). Duplicates were averaged to determine the average *T_m_* of all identified venom toxins. Venom proteins failing to reach 50% denaturation even at higher temperatures were classified as non-melting proteins.

### Analysis of PISA data

PISA data was analyzed as previously described (71,98). Briefly, PISA uses the ΔS_m_ value or soluble abundance ratio (SAR) as opposed to the *T_m_* to determine differences in thermal stability (70). ΔS_m_ represents the difference in integral abundance of a protein in treated compared to untreated samples. We performed a two-tailed Student’s *t*-test with unequal variance to calculate p-values (p<0.05). Changes to venom protein abundance were visualized using volcano plots based on log_2_SAR values and -log_10_ transformed p-values. Proteins with a log_2_SAR value ≥ 0.5 and a -log_10_ transformed p-value ≥ 1.3 (p<0.05) were identified as toxins with a significant shift to thermal stability. All figures were made with BioRender.com.

## Results

### Crotalus atrox venom proteome

Previous characterization of the *C. atrox* venom proteome revealed the presence of at least 24 proteins belonging to eight different venom toxin protein families (Figure 2a; (95)) and, more recently, the presence of 31 SVMP genes in *C. atrox* with 15 to 16 expressed SVMPs (100). SVMPs and snake venom serine proteases (SVSPs) were the two most abundant protein families representing nearly 70% of the venom proteome. L-amino acid oxidase (L-AAO), Phospholipase A_2_ (PLA_2_), disintegrins, and cysteine-rich secretory proteins (CRISPs) comprise most of the remaining 25% of *C. atrox* venom proteins, whereas vasoactive peptides, endogenous SVMP inhibitors, and C-type lectins (CTL) comprised the remaining small fraction of venom components comprising <2% of the venom proteome. Utilizing the protein databased generated using sequences of proteins identified by Calvete et al. (95) combined with protein sequences derived from a *C. atrox* venom gland transcriptome, we detected 46 unique proteoforms with at least one unique peptide in *C. atrox* venom (Figure 2b). Venom toxins with the highest number of distinct proteoforms detected included 13 CTLs, 13 SVMPs, nine SVSPs, and three PLA_2_s (Figure 2b). We identified only one unique proteoform of more abundant proteins including L-AAO and CRISP and only one proteoform for minor components bradykinin-potentiating peptide (BPP), glutaminyl-peptide cyclotransferase (GPC), hyaluronidase (HYAL), nerve growth factor (NGF), phospholipase B (PLB), and vascular endothelial growth factor (VEGF).

**Figure 2.**
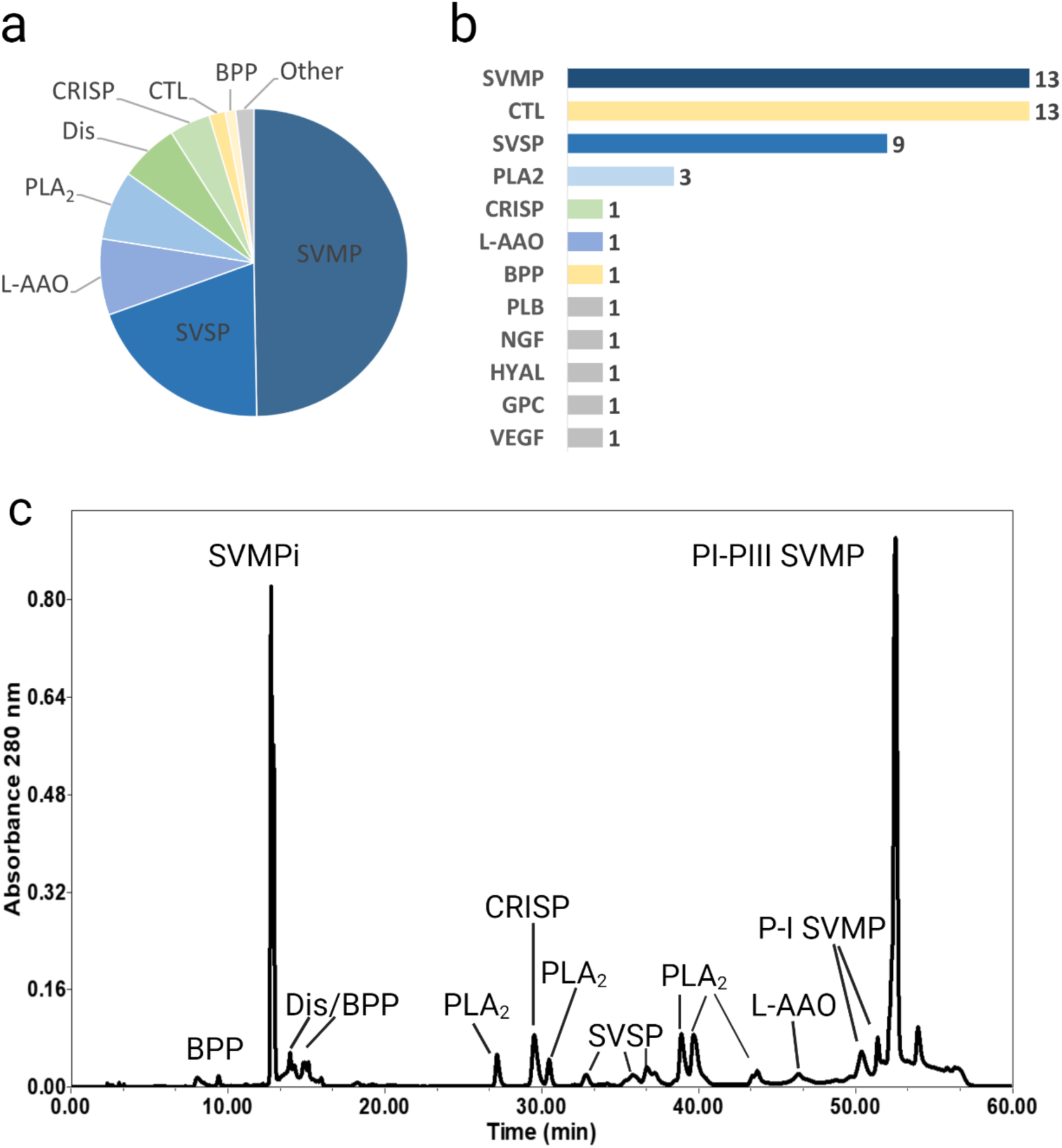
*Crotalus atrox* venom proteome characterization. a) Toxin family abundances in *C. atrox* venom modified from Calvete et al., 2009. b) The number of proteoforms identified in *C. atrox* venom in the present study organized by family. c) RP-HPLC separated *C. atrox* venom. For peak identification fractions were analyzed with mass spectrometry and SDS-PAGE and compared to known masses from Calvete et al., 2009. SVMP=Snake venom metalloprotease, CTL=C-type lectin, SVSP=snake venom serine protease, Dis=disintegrin, PLA_2_ =phospholipase A_2_, BPP=Bradykinin potentiating peptide, CRISP=cysteine rich secretory protein, L-AAO=L-amino acid oxidase, SVMPi= SVMP tripeptide inhibitor, PLB= Phospholipase B, NGF=nerve growth factor, HYAL=hyaluronidase, GPC=glutaminyl-peptide cyclotransferase, VEGF=vascular endothelial growth factor.

RP-HPLC analysis revealed a complex toxin profile of *C. atrox* venom similar to that of Calvete et al., 2009 (Figure 2c). We used SDS-PAGE of peak fractions (Supplemental Figure 1) combined with the peak elution times and identified masses in Calvete et al. (95) to confirm peak identities. BPP’s eluted between 8 and 15 min with co-elution of disintegrins and SVMP inhibitors. PLA_2_ eluted at 27, 30, and between 39-44 min, CRISP eluted at 29 min, SVSP eluted between 32 – 36 min, and L-AAO eluted at 46 min. SVMPs eluted between 51-56 min (Figure 2c).

### C. atrox venom meltome

With the goal of demonstrating the utility of applying a TPP workflow for identifying venom protein interactions with a small molecule inhibitor, we first assessed the effects of thermal stress on the venom proteome. Venom was subjected to increasing temperatures ranging from 40 to 75°C, allowed to cool at room temperature, followed by separation and removal of aggregates from each temperature point by centrifugation. The soluble fractions were then visualized by gel electrophoresis (Figure 3a) and prepared for LC-MS/MS. SDS-PAGE analyses of these fractions indicate that the entire venom proteome appeared to exhibit some degree of denaturation between the temperatures tested with clear differences in denaturation observed across venom protein families (Figure 3a). For example, L-AAO, HYAL, SVMPs, and CTLs appeared more thermally sensitive while SVSP, CRISP, PLA_2_, and disintegrin families exhibited greater stability at higher temperatures.

**Figure 3.**
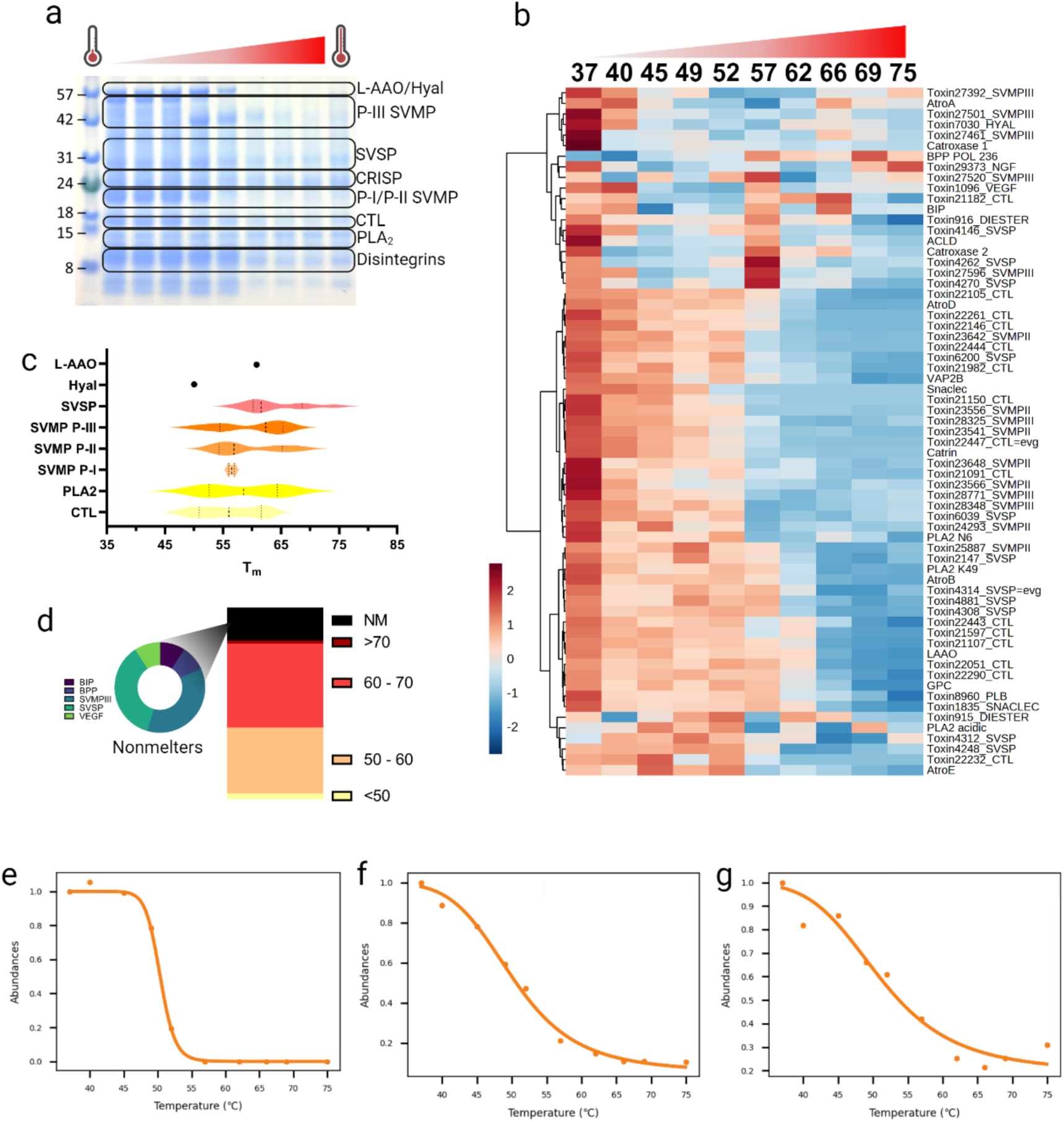
*C. atrox* meltome characterization. a) SDS-PAGE of *C. atrox* venom heated at temperatures between 37-70°C for 3 minutes. b) Heatmap showing the thermal denaturation of most toxin proteoforms across 10 temperatures where 37°C represents the nondenatured control. Heatmap colors represent normalized intensity and are scaled by row to better visualize variation in intensity between temperatures. c) Distribution of melting temperatures organized by family. Dotted lines represent median and quartile ranges. d) Distribution of melting temperatures for all toxins identified. Nonmelters (NM) are classified as proteoforms for which *T_m_* could not be calculated when heated to a maximum temperature of 75°C. e) Representative melting curve of a CTL (Crotocetin). Abundance is normalized to 37°C f) Representative melting curve of a PLA_2_ (Cvv-N6). Abundance is normalized to 37°C. g) Representative melting curve of an SVMP (PIII 28325). Abundance is normalized to 37°C.

Next, we assessed the thermal stability of venom proteins across the 10 different temperature points by LC-MS/MS. An equal volume of each soluble fraction was collected and subjected to reduction, alkylation, trypsin digestion, and LC-MS/MS. Fragmentation spectra were interpreted against our *C. atrox-*specific custom venom proteome sequence database, and we used the ProSAP package (99) to determine melting points for each venom protein family. When normalized to thermostable proteins, most venom proteins show decreasing abundance with increasing temperature, with the majority of proteins reduced in abundance at temperatures above 62°C (Figure 3b). The distribution of toxin melting temperature (*T_m_*) values ranged from 47.8-74.3°C (Figure 3c and 3d). Most toxins had *T_m_*’s between 50-60°C (Fig 3d; n=22) or 60-70°C (n=28), and only 11 proteoforms were still thermostable with no calculable *T_m_* at 75°C (BIP, BPP, VEGF, 4 SVMPIII, and 4 SVSPs; Fig 3d). The five toxins with the lowest *T_m_*’s included four CTLs (average *T_m_* =49.3°C, stdev=1.1°C; Figure 3c and 3e) and the single hyaluronidase proteoform (*T_m_* =50.6°C). CTLs *T_m_* values as a whole ranged from 47.8-63.1°C (ave= 56.0°C, stdev=5.3°C). PLA_2_s had an average *T_m_* of 61.2°C (stdev=8.3°C; Figure 3c and 3f). The different SVMP subfamilies differed slightly in their melting range but were not significantly different (p=0.77; Figure 3c). PI-SVMP proteoforms had an average *T_m_* of 56.8°C (stdev=0.03°C), PIIs averaged 59.3°C (stdev=5.7°C; Figure 3c), and PIII’s averaged 59.7°C (stdev=6°C; Figure 3g). SVSPs had the highest average *T_m_* (63.9°C, stdev=5.5°C), and made up a large proportion of the proteins that were thermostable above 75°C (Figure 3b). The single L-AAO proteoform identified melted at 61.7°C. In general, melting temperatures were reproducible between replicates with an average standard deviation of 1.17°C between replicates. These results demonstrate protein family-level differences in thermal stability, in that all proteoforms of some families denatured (i.e., CTL, SVMP I) when subjected to heat, while others appear resistant to thermal perturbation (SVSPs). These results indicate that a significant proportion of the venom proteome is amenable to thermal denaturation.

### Venom-wide interactions with marimastat

After establishing that venom proteins are susceptible to thermal denaturation, we next assessed if a TPP strategy could be applied to elucidate small molecule-venom protein engagement. For this, we applied the PISA assay, a simplified TPP approach where samples across the entire temperature gradient of the same treatment are pooled prior to preparation and mass spectrometric analysis (66,70,71). With traditional PISA, the abundance of the protein(s) in the soluble fractions of the pooled samples is then used to assess the effect of a compound on its thermal stability (70). For highly thermostable proteins, monitoring supernatant alone is likely not effective in thermal-shift-based methods (76). Because binding of a compound can, in some cases, lead to protein destabilization rather than stabilization, quantifying protein abundance in the precipitate pellet can also identify protein targets (76). Further, because of different observed signal-to-noise ratios, soluble and pelleted material may perform differently in PISA assays to identify significant thermal shifts (75,76). Utilizing precipitated material to measure changes in protein stability can additionally reduce the false discovery rate (FDR) and improve sensitivity of the assay. Based on this logic, we utilized both supernatant and precipitated material to investigate the effects of marimastat.

PISA assays were performed on marimastat, an inhibitor of matrix metalloproteases that has shown significant inhibitory activity against SVMPs (50,51,53,56,60–62). *Crotalus atrox* venom was incubated with marimastat (15 μM and 150 μM) or vehicle (ddH_2_O) for 30 min at 37°C. Following incubation, each sample was divided into 12 aliquots and subjected to increasing temperatures from 40 to 70°C. Equal aliquots per temperature point were then pooled, protein aggregates were separated by centrifugation, and the soluble and insoluble fractions of the vehicle and inhibitor-treated venoms were prepared for downstream analysis.

When filtering criteria were applied (p<0.05, and log_2_SAR >0.5), the lower concentration of marimastat (15 μM) caused five of 21 SVMP proteoforms supernatant (PIII 28348, PIII 28325, PII 25887, PII 23541, and VAP 1) to display a stabilizing shift in treated supernatant compared to untreated supernatant (Figure 4a). In addition to these five proteoforms, two additional SVMPs (PII 23556 and PII 27392) were more abundant in the supernatant of venom treated with 150 μM of marimastat (Figure 4b).

**Figure 4.**
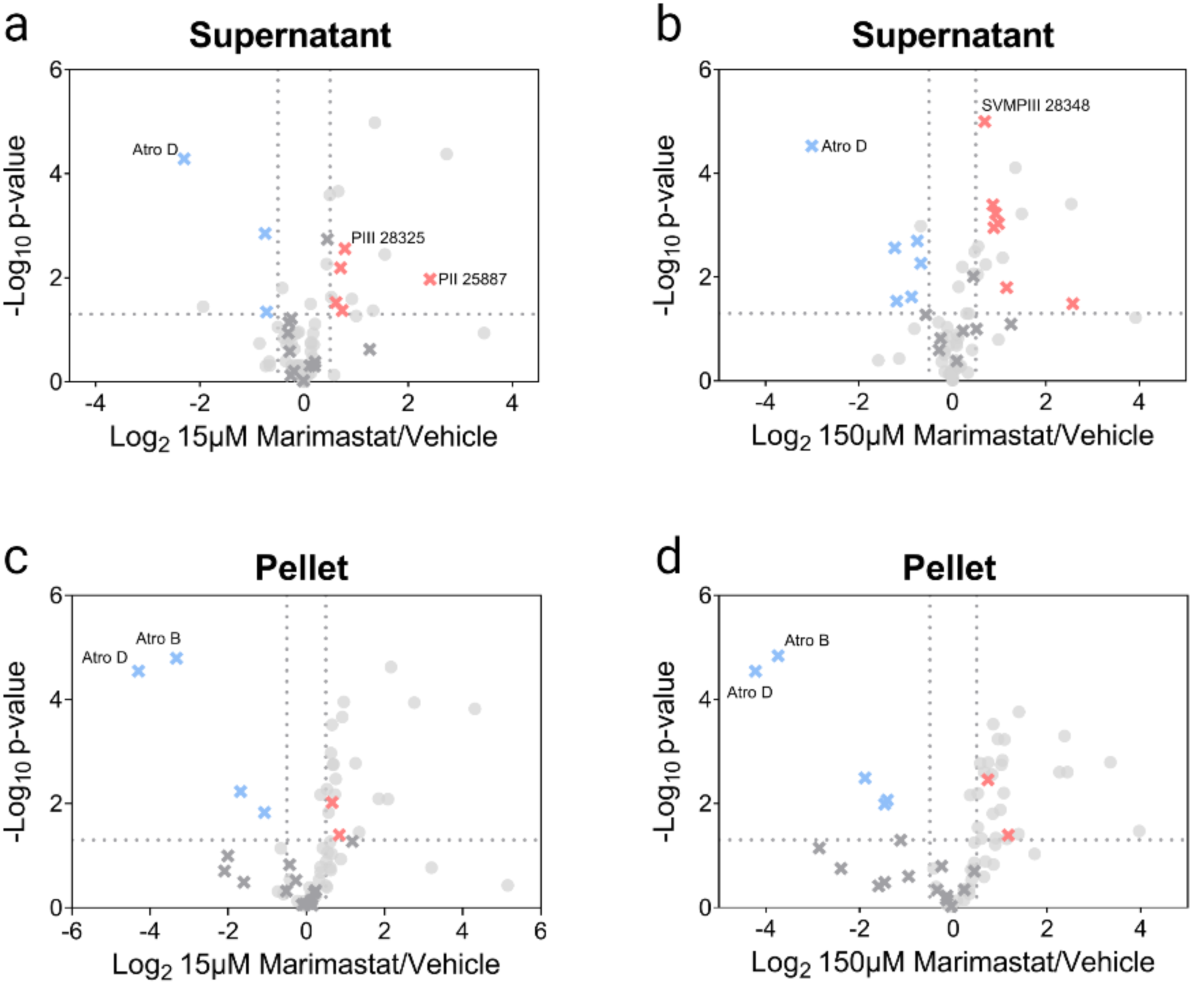
*C. atrox* venom-wide interactions with two concentrations of marimastat with temperature window from 40-70°C. a) Volcano plot comparing soluble supernatant of heat-treated venom + marimastat (15 μM) to heat-treated venom alone. X indicates SVMP proteoforms, red=positive outliers, blue=negative outliers, grey=not significant. b) Volcano plot comparing soluble supernatant of heat-treated venom + marimastat (150 μM) to heat-treated venom alone. c) Volcano plot comparing insoluble precipitate of heat-treated venom + marimastat (15 μM) to heat-treated venom alone. d) Volcano plot comparing insoluble precipitate of heat-treated venom + marimastat (150 μM) to heat-treated venom alone.

Next, we compared the pellets of untreated venom to venom treated with both concentrations of marimastat. When filtering criteria were applied at the low concentration, only four SVMPs (Atro B, Atro-D, SVMPIII 27520, and SVMPIII 28348) were detected at significantly lower abundance in the treated pellet compared to the control pellet, indicative of a stabilizing effect of marimastat (Figure 4c). These same proteoforms, in addition to SVMP PII 23541, were also significantly reduced in the pellet of the higher marimastat concentration (Figure 4d). The presence of positive outliers identified in both the supernatant and negative outliers in the precipitate of treated venom indicates an overall stabilizing effect of marimastat on venom targets.

### Validation of inhibitor interactions

To validate our PISA results showing the stabilizing effects of marimastat on SVMPs, we performed SDS-PAGE and RP-HPLC on non-heat-denatured venom and venoms treated with marimastat or vehicle. The toxin family composition of the peaks and bands altered by the addition of marimastat was confirmed by mass spectrometry (Supplemental Tables 1-2). In the heated marimastat-treated venom, gel bands A and B were composed predominantly of PIII 27501 and VAP2B (band A) and PII 23556, Atro E and Atro B (band B; Figure 5a). Band C was composed of acidic PLA_2_, band D of PLA2 Cax-K49, CTL 22443, and PLA2 Cvv-N6, and band E was predominantly CTL 21182, CTL 22444, and PLA2 Cax-K49. SDS-PAGE analysis of the heat-denatured and undenatured control venom shows a clear reduction in the size and intensity of SVMP-PIII (∼50kDa; band A), SVMP P-I/II (∼20kDa; band B), and CTL/PLA_2_ gel bands (∼10-14kDa; bands C-E) in response to thermal treatment (Figure 5a). This reduction in SVMP and CTL/PLA_2_ band size and intensity in response to heat appears to be partially to fully recovered when venom is incubated with 150 μM marimastat (Fig. 5a).

**Figure 5.**
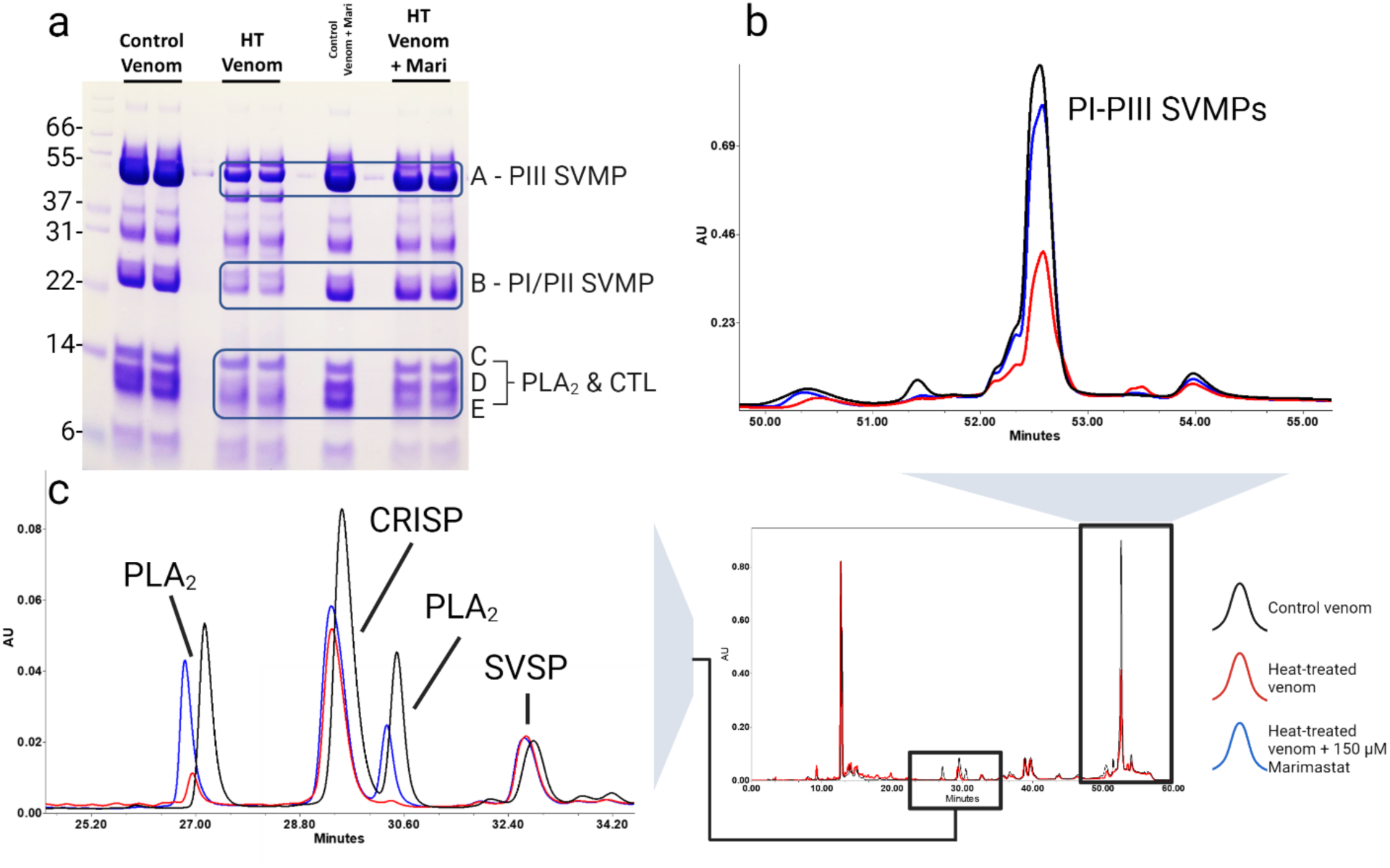
Validation assays of inhibitor interactions. a) SDS-PAGE comparison of heat-treated venom to heat-treated venoms incubated with 150 μM of marimastat with a thermal window of 40-70°C. Note the recovery in band size and intensity of SVMP and PLA_2_ bands in marimastat-treated samples. Left side indicates molecular mass standards in kDa. b) Enlarged HPLC-separated SVMP peak overlay comparing abundance of non-heat-treated venom (black), heat-treated venom (red), and venom heat-treated after incubation with 150 μM of marimastat (blue). Note the recovery of peak area in the inhibitor-treated sample. c) Enlarged HPLC-separated PLA_2_ peak overlay comparing abundance of non-heat-treated venom (black), heat-treated venom (red), and venom heat-treated after incubation with 150 μM of marimastat (blue). Note the recovery of PLA_2_ peak area in the inhibitor-treated sample.

RP-HPLC peaks eluting between 51 to 54 minutes were identified by mass spectrometry as SVMPs, with VAP2B, PIII 27501, P-III ACLD, PII 23556, PIII 28348, PII 23566, and Atro E representing the dominant proteoforms. These continued to be the dominant proteoforms with the exception of Atro E in both the heated control and marimastat-treated venoms. The 27-minute, 29-minute, and 30-minute peaks were composed predominantly of the basic PLA_2_ Cax-K49, CRISP, and basic PLA_2_ Cvv-N6 respectively (Figure 5c). These remained the dominant proteoforms in the heated control venom and marimastat-treated venom, with the exception of marimastat-treated peak 30 where CRISP became the dominant proteoform followed by PLA_2_ Cvv-N6.

The stabilizing effect of marimastat on some venom proteins is further demonstrated by analysis of RP-HPLC, which shows partial recovery in the chromatographic peak area and peak height of SVMP and two PLA_2_ peaks in the marimastat-treated venoms compared to the controls (Fig 5b-c). After heat treatment, SVMPs lose 44% of their original peak area, but marimastat treatment results in only a 7% decrease in peak area after melting (Figure 5b). The PLA_2_ proteins eluting at 27 minutes decreases by 75% when venom is heat-treated and only 26% when venom is treated with marimastat, while the PLA_2_ eluting at 30 minutes is virtually absent in the heated control venom but only loses 52% abundance when heat-treated with marimastat (Figure 5c).

Peak heights of PLA_2_ (27 minutes), PLA_2_ (30 minutes), and SVMPs decrease by 81%, 100%, and 57% respectively after melting; however, with marimastat peak height only decreases by 21%, 48%, and 21% respectively (Figure 5b-c). VAP2B is the dominant proteoform in *C. atrox* venom (Figure 5b; Supplemental Table 2) and was the second most abundant proteoform in the SVMP fractions and gel bands of treated venom. However, it was not detected as a stabilized outlier in either supernatant or pellet in the current PISA experiments performed with the temperate range of 40 to 70°C. Thus, we aimed to increase the sensitivity of the PISA assay with a narrower thermal window determined by the T_m_ values previously calculated for the target toxin family.

### Venom-wide interactions with marimastat in a narrowed thermal window PISA

While PISA is advantageous because it reduces the analysis time and sample preparation while still being effective at target discovery, it may sacrifice sensitivity compared to TPP experiments due to the pooling of all temperature points, and melting temperature selection can have a drastic effect on thermal behavior in PISA experiments (73,76). To investigate this, we compared the performance of a broad thermal window (40-70°C) to a narrower window (56-60°C), selected based on the mean and standard deviation of *T_m_*’s of the target venom toxin family. We performed PISA assays with the same concentrations of marimastat, with a narrower temperature window from 56-60°C with 5 temperature points of each sample replicate, which has been shown to improve the overall sensitivity of the PISA assay in target identification (73).

When samples were heat treated with a narrower window of temperatures, the lower concentration of marimastat displayed three of 21 SVMP proteoforms (PIII 28348, PIII 28325, PII 23541) at a greater abundance in treated supernatant compared to untreated supernatant (Figure 6a). At the higher concentration, seven of 21 proteoforms were higher in supernatant of treated venom: VAP1, PIII 28348, PIII 28325, PII 23556, PII 23541, PIII 28771, PIII 27392 (Figure 6b). In the pellets of samples treated with a narrower range of temperatures, 11 of 21 proteoforms (VAP2B, PIII 28348, Atro D, PIII 27501, PII 23541, Atro B, PIII 27461, PIII 27520, PIII 28771, PII 23648, PIII 27392) demonstrated a stabilizing shift in the pellet of the lower concentration of marimastat condition (Figure 6c). These same proteoforms plus PII 23556 were reduced in the pellet at the higher concentration of marimastat (Figure 6d).

**Figure 6.**
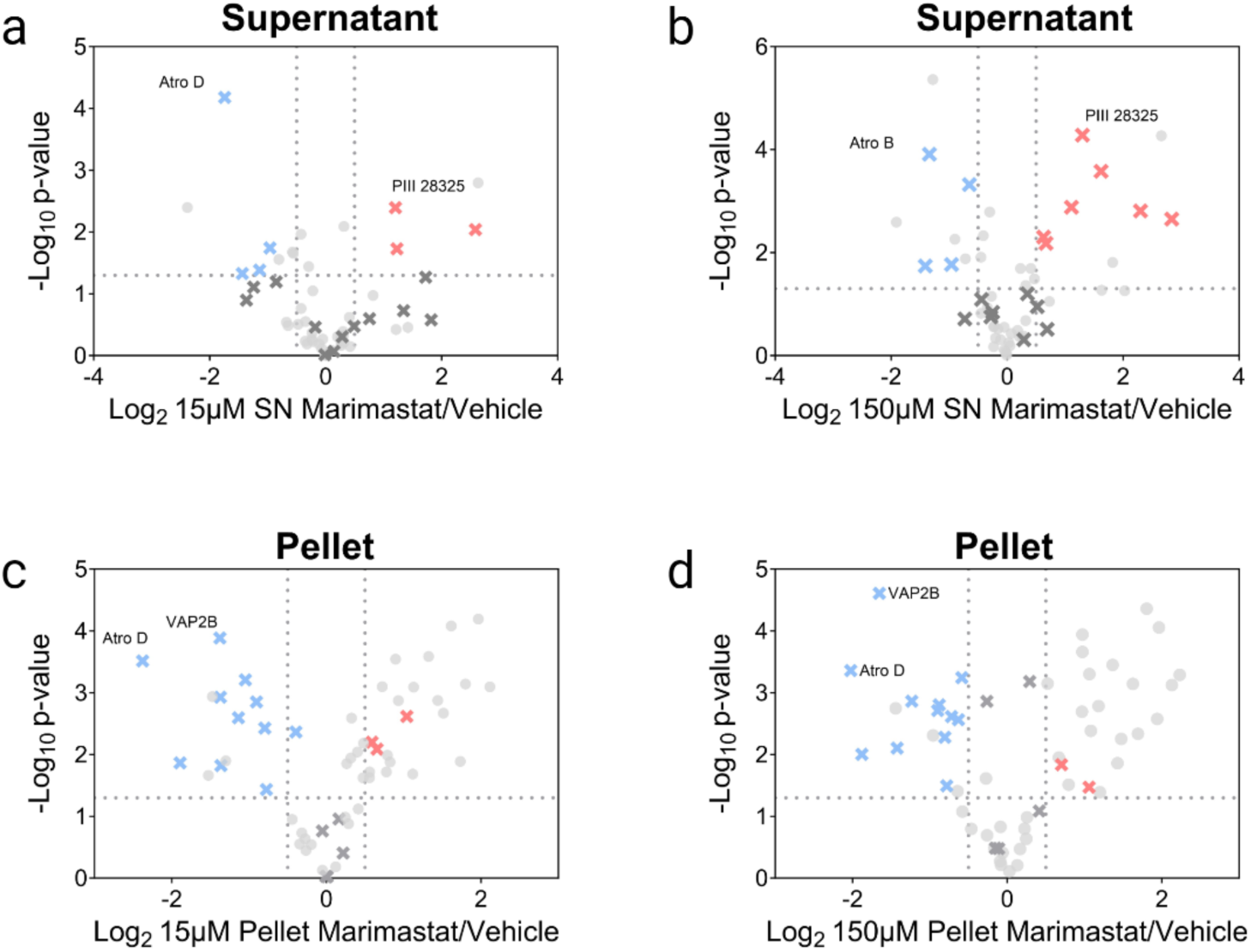
*C. atrox* venom-wide interactions with two concentrations of marimastat with temperature window from 56-60°C. a) Volcano plot comparing soluble supernatant of heat-treated venom + marimastat (15 μM) to heat-treated venom alone. X indicates SVMP proteoforms, red=positive outliers, blue=negative outliers, grey=not significant. b) Volcano plot comparing soluble supernatant of heat-treated venom + marimastat (150 μM) to heat-treated venom alone. c) Volcano plot comparing insoluble precipitate of heat-treated venom + marimastat (15 μM) to heat-treated venom alone. d) Volcano plot comparing insoluble precipitate of heat-treated venom + marimastat (150 μM) to heat-treated venom alone.

### Heat-treatment comparison

Next, we compared the performance of a broad thermal window (40-70°C) and narrower thermal window (56-60°C) to our results gathered from our validation experiments (Figures 4-6). At both temperature ranges when venom was treated with marimastat, principal component analysis (PCA) shows clustering of the replicates based on treatment condition, with the two marimastat-treated groups separating from the vehicle-treated samples (Figure 7a-d). These results indicate that both concentrations of marimastat interact with venom protein targets and alter the thermal stability of venom proteins compared to the control group. However, pellet replicates (Figures 7c-d) cluster more tightly together in both conditions than in supernatants with greater separation among the treatment groups (Figures 7a-b). The highest amount of variance explained (97%) by the top two principal components was in the narrow window pellet, though all plots had a high percentage of sample variance explained (>86%). The number of significantly stabilized proteins found in the supernatant (p<0.05, log_2_SAR >0.5) after treatment with marimastat at a broad melting window were 12 and 14 for 15 μM and 150 μM, respectively (Figure 7e). The percentage of SVMPs among the identified proteins were 42% and 50%. At the narrower melting window, four and nine proteins were identified in 15 μM and 150 μM treatments, respectively, but SVMPs comprised 75% and 78% of identified proteins. In general, pellets of both melting windows appeared to perform better regardless of concentration. The precipitated pellet from 150 μM-treated venom heated at the narrower thermal window identified the most SVMP proteoforms of any treatment group. Though the broader temperature window precipitate identified fewer SVMP proteoforms, SVMPs were the only venom toxin family proteoforms identified, while the supernatant appeared to contain more, potentially off-target, non-SVMP identifications. The broad melting window identified four and five SVMP proteoforms, while the narrow window pellets identified 11 and 12 for 15 μM and 150 μM, respectively (Figure 7e).

**Figure 7.**
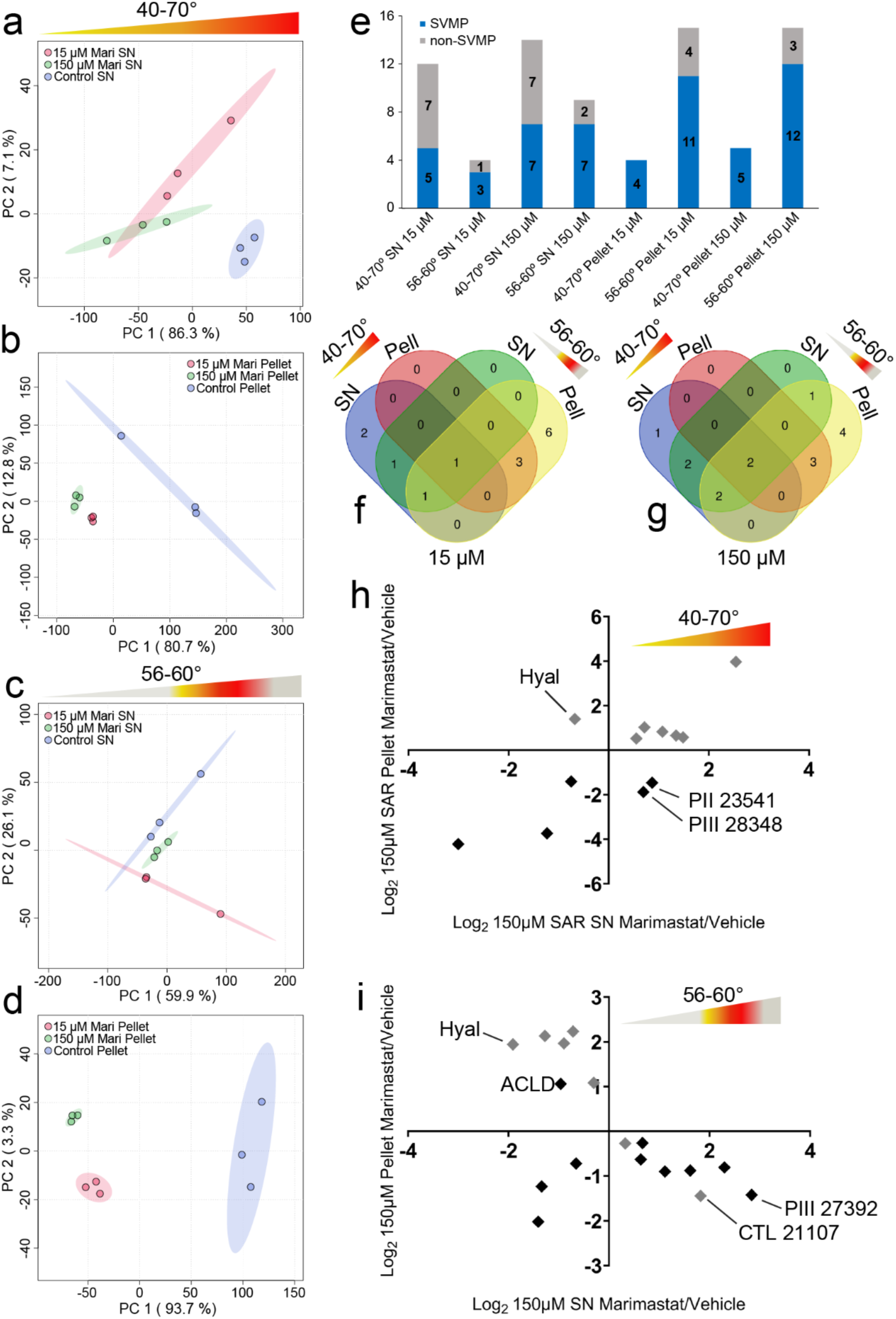
Comparison of a broad (40-70°C) to a narrow (56-60°C) PISA thermal window. PCA plot comparing replicates of soluble supernatant of a) heat-treated venom + 15 μM marimastat to heat-treated venom alone and b) heat-treated venom + 150 μM marimastat to heat-treated venom alone. SN= supernatant, Pell= pellet, Con= control. PCA plot with 95% confidence intervals comparing replicates of insoluble precipitate of c) heat-treated venom + 15 μM marimastat to heat-treated venom alone and d) heat-treated venom + 150 μM marimastat to heat-treated venom alone. SN= supernatant, Pell=pellet. SN= supernatant, Pell= pellet, Con= control. e) Number of SVMP and non-SVMP proteins identified with significant thermal shifts toward stabilization (p-value<0.05 log_2_SAR>0.5) after 15 μM or 150 μM marimastat treatment in supernatants and pellets heat-treated at a broad (40-70°C) or a narrow (56-60°C) thermal window. SN=supernatant, P=pellet. Venn diagrams of SVMP proteins identified with significant thermal shifts toward stabilization (p-value<0.05, log_2_SAR>0.5) in supernatants and pellets heat-treated at a broad (40-70°C) or a narrow (56-60°C) thermal window after f) 15 μM or g) 150 μM marimastat treatment. SN= supernatant, Pell=pellet. Scatter plot showing log_2_SAR values calculated from 150 μM marimastat-treated venom vs. vehicle treatment heated at from h) 40- 70°C or i) 56-60°C that meet significance criteria in both supernatant (SN) and precipitate (Pellet) group. Largest outliers of SVMP and non-SVMP proteoforms are labeled. SN=supernatant. Black=SVMP proteoforms, grey=non-SVMP toxins. SN= supernatant, Hyal=hyaluronidase, ACLD=PIII SVMP ACLD, CTL= C-type lectin.

Performance of specific SVMP proteoform identification between melting windows varied significantly at both concentrations, with only one common proteoform at 15 μM (Figure 7f) and two at 150 μM (Figure 7g). At both concentrations, the narrow window pellets had the highest number of uniquely identified SVMP proteoforms. When soluble abundance ratios (log_2_SAR) of supernatant and pellets are compared, the narrower thermal window performs significantly better at identifying target and off-target proteoforms that meet significance criteria in both supernatant and pellet. When only proteins meeting significance criteria for both supernatant and pellet were compared, the broad window performed poorly, identifying only two SVMPs (PIII 28348, PII 23541). with significant stabilizing shifts (Figure 7h). The narrower window identified six SVMP proteoforms with significant stabilizing shifts (VAP1, PII 23556, PIII 28348, PIII 28771, PII 23541, PIII 27392) and one with a possible destabilizing shift (ACLD; Figure 7i). Off-target proteins that met significance criteria for both conditions included hyaluronidase, and, using the narrower window, three SVSPs and VEGF.

### SVMP comparisons and target identification

Because the narrower temperature window appears to identify more target proteoforms with less noise, we utilized this narrow window approach to re-analyze interactions between SVMPs and marimastat. More SVMP proteoforms were identified as significant with a narrower window when analyses of the supernatant and pellet are combined (Figure 7e-g), and analyses of the pellet identified more SVMP proteoforms than the supernatant within the narrower thermal window (Figure 8a). Specifically, analysis of the narrow range pellet identified highly abundant proteins also identified in the validation assays but not identified using the broad the broad thermal window (e.g., VAP2B). Hierarchical clustering analysis comparing supernatants and pellets at both concentrations shows an inverse relationship between relative intensity of each proteoform in the pellet vs. the supernatant (Figure 8b). When these conditions are compared to controls, we resolved three patterns of various proteoforms: 1) proteoforms that showed a positive shift (trend towards stabilization) in the supernatant at both concentrations of marimastat; 2) proteoforms that disappear from the pellet after marimastat treatment but do not necessarily increase in SAR in the supernatant (trend towards stabilization); and 3) proteoforms that increase in pellet SAR after treatment (Figure 8c). Finally, correlation analysis performed with the SAR values of proteoforms supernatants and pellets of the narrow thermal window identified three clusters of proteoforms with similar shifts in thermal behavior: 1) a cluster containing the proteoforms that were stabilized by marimastat (e.g., VAP2B, 27501, 23556, 27392) that represent the strongest targets of marimastat, 2) a cluster containing atrolysin B, atrolysin D, and PIII 27520 which appeared to decrease in abundance in both supernatants and pellets after marimastat treatment, and 3) a cluster that did not appear to be thermally stabilized by marimastat including atrolysin A, PII 24293 (Figure 8d). The strongest target list includes proteoforms identified in the validation protein gel (e.g., VAP2B, PIII 27501, PII 23556, PIII 28348, PIII 27392, and PII 23648), and those identified as most abundant in the SVMP HPLC peaks (VAP2B, PIII 27501, PII 23556, PIII 28348, PIII 27461).

**Figure 8.**
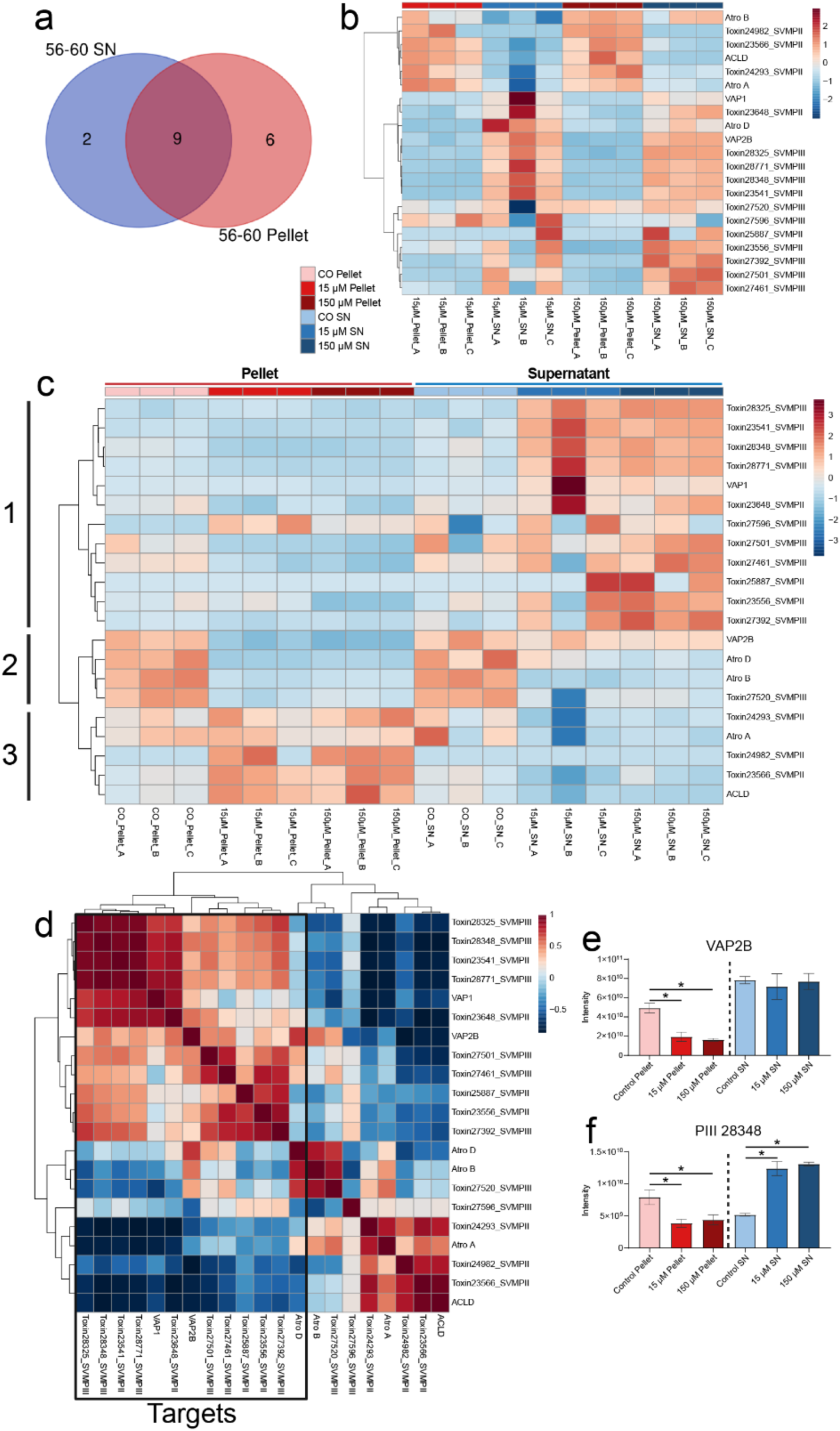
Effects of marimastat treatment and a narrow thermal window on SVMP proteoforms only. a) Number of proteins identified in supernatant (SN) and pellet of narrow thermal window that meet significance criteria (p-value<0.01, log_2_SAR>0.5) b) heatmap of sum-normalized intensity values in supernatant or precipitate of SVMPs from 15 μM or 150 μM marimastat treated venom. Heatmap colors are scaled by row to better visualize variation in sum-normalized intensity between classes. c) heatmap of sum-normalized intensity values in supernatant or precipitate of SVMPs from 15 μM or 150 μM marimastat treated venom or vehicle control showing concentration-dependent shifts in abundance. Heatmap colors are scaled by row to better visualize variation in sum-normalized intensity between classes. d) Correlation plot of SAR values showing strongest marimastat targets based on effects of marimastat treatment on SVMP proteoform intensity. Comparison of concentration-dependent intensity shifts between precipitate and supernatant of e) the most abundant SVMP proteoform VAP2B and f) a less abundant SVMP proteoform PIII 28348 at both concentrations of marimastat at the narrow thermal window.

The most abundant proteoforms were identified as significant in analysis of the pellet but not in the corresponding supernatant. To further explore the apparently superior performance of analysis of the pellet for identifying high abundance targets, we use VAP2B as an example. In the case of VAP2B, the stabilizing effect of marimastat is only evident in the pellet, and both 15 μM and 150 μM concentrations have significantly lower levels of the precipitated toxin (p=0.003, p=0.0013, respectively, Figure 8e). A less abundant proteoform, PIII 28348, had a stabilizing shift that was detected in both the pellet and the supernatant at both concentrations of marimastat (Figure 8f). In the pellet, abundance of PIII 28348 was significantly lower at both concentrations than in the untreated control (p=0.0004, p=0.0014, respectively), and significantly higher in the supernatant compared to control (p<0.0001).

### Off-target effects

Based on recovery of peak area in PLA_2_ and CTL-containing peaks and gel bands in marimastat-treated venom subjected to RP-HPLC (Figure 5c), we explored the possibility of using PISA assays to detect off-target effects of marimastat. Interestingly, at both concentrations with the wider thermal window we observed changes in thermal behavior indicative of a shift towards stabilization of some non-target protein families, including two PLA_2_ proteoforms (Cax-K49 and Cvv-N6) and four CTL proteoforms. In the gels, there was recovery of band E containing Cax-K49, and CTL’s 22443, 21182, and 22444. Both PLA_2_ proteoforms were repeated positive outliers in supernatants of all conditions but were not significant in any pellets. Various CTL proteoforms including CTL 21107, 22105, 22447, 21150, and 22232 were significant outliers in some conditions. When comparing only significant log_2_SAR values in both supernatant and pellet, hyaluronidase, VEGF, 2 SVSPs and 2 CTL’s (22444, 21107) were significantly correlated between pellet and supernatant at the narrower melt window, but only hyaluronidase displayed stabilizing behavior (Figure 7i).

## Discussion

The development and testing of alternative snakebite therapeutics that are affordable, stable, and easily administered is an urgent global need (1,10,47,101). Small molecule inhibitors currently lead the field of possible supplementary snake envenomation therapies, with phase II clinical trials ongoing for the PLA_2_ inhibitor varespladib (102) and the SVMP inhibitor DMPS ((103); Clinical Trials.gov, 2021). Numerous inhibitors have shown promising cross-species efficacy *in vivo* and *in vitro* (37,50,53,57,104), indicating that they may be less vulnerable to the effects of venom variation than traditional antibody-based antivenoms. However, additional preclinical studies are needed to evaluate the neutralizing efficacy and specificity of these drugs alone and in combination, and the development of these drugs would be accelerated by implementation of high throughput screening of interactions and efficacy across many species. Research on small molecule inhibitors of snake venom toxins has typically focused on *in vitro* and *in vivo* functional assays based on the known or likely biological activities of toxins (49,50,108,51–53,58,104–107). These approaches utilize a downstream measurement of the presumed interactions of an inhibitor with its targets (ex. reduced specific activity or increased survival). A previous study performed molecular docking analysis using marimastat and a purified PI SVMP proteoform CAMP-2 to demonstrate a direct interaction (51). However, the PISA method outlined here represents both a direct and venom-wide assessment of target-ligand engagement and provides the opportunity to link direct target-ligand interactions with functional and phenotypic responses (71,72,109).

In this study, we investigate the thermal characteristics of the *C. atrox* venom proteome and use this to develop a PISA-based assessment of the venom proteome-wide targets of the SVMP inhibitor marimastat. We investigate both its proteome-wide effects and determine and validate its interactions with specific venom proteoforms of its target toxin family (SVMPs) as well as possible off-target protein families. We identified a suite of marimastat proteoform-level targets and confirmed them by RP-HPLC and SDS-PAGE. We also compared the performance of soluble supernatant and insoluble precipitate at two different inhibitor concentrations for target identification. Our results provide a promising first assessment of the application of a PISA-based approach as a sensitive and high-throughput method to assess the direct targets of small molecule inhibitors for snake venom. Based on our experiments with PISA in this context, we find that analysis of the insoluble fraction from venom that was treated with a high concentration of marimastat, but a narrow thermal window for PISA, provided more sensitive target data with the least noise.

Previous research has shown that small molecule inhibitor efficacy *in vitro* may not always translate to *in vivo* efficacy. For example, the SVMP inhibitors dimercaprol and prinomastat showed moderate to high SVMP inhibitory activity *in vitro* but failed to confer any protection towards crude venom in *in vivo* assays (60). Furthermore, studies have highlighted cross-species variation in neutralization effects of potential inhibitors, which has significant implications for the application of inhibitors as broadly effective pre-hospital treatments of envenomation by potentially diverse species (58). Dimercaprol showed promise in murine models as an SVMP inhibitor against *Echis ocellatus* venom (110), however, it lacked this protective effect *in vivo* against *Dispholidus typus* venom (60), likely due to the high levels of divergence in venom composition between these distantly related species. While some inhibitors have demonstrated neutralization capacity of specific biological effects (such as anticoagulation) caused by venoms of divergent species, they may vary in effectiveness across species because of lineage-specific variation in venom toxin sequence, activity, or relative abundance or because the same biological effects may arise due to the action of different toxin families altogether (58,60). Knowledge of the snake species-specific venom-wide and proteoform-specific efficacy of inhibitors has the potential to significantly improve our ability to predict cross-species neutralization and to unravel the disparity between *in vitro* and *in vivo* results.

Before PISA could be widely applied for the screening of a large number of potential inhibitors against snake venom, a number of considerations must be addressed. By pooling a wide range of temperature points, PISA data in particular may suffer from reduced screening sensitivity, depending on the specific thermal properties of various proteins (73). Venom toxins appears to have significantly higher T_m_ values than human cell types, which ranged from 48 to 52°C (92). Some potential snake venom toxin families of interest (i.e., SVSPs and CRISPs) display high thermal tolerance, which generally suggests that a thermal shift assay would be less than ideal to investigate inhibitor-toxin interactions for such thermostable proteoforms. Based on the target family-level thermal properties determined by TPP, we refined our PISA assay parameters to a more sensitive thermal window for target identification and showed that a narrower thermal window selection can improve inhibitor target identification. These findings highlight how knowledge of general thermal properties of a toxin family of interest might be used to improve target identification, perhaps even for protein families with higher thermal stability.

Our results demonstrate how analysis of the composition of both supernatants and pellets can be complementary, and thus be integrated to further refine inferences of molecular targets (75,76). In our experiments, we observed varying performances between supernatant and pellet data in the consistent identification of inhibitor targets, particularly of the high abundance SVMP proteoform VAP2B. We found that precipitated material of the narrowed thermal window provided enhanced sensitivity for target deconvolution of the most abundant toxins and across the proteome in general. As previously noted, precipitated material produces better signal-to-noise ratios and more apparent stability ratios compared to analysis of supernatant (75). Indeed, some previously investigated well-known drug targets were only identified in the precipitated material, with no corresponding stability ratio shift in the supernatant (75), as seen with VAP2B in our study, indicating that pelleted material is not just complementary to supernatant-based results, but may be critical for thorough target deconvolution. This is likely due to the continued presence of many proteins even at high temperatures as observed in this study and in previous studies (75). We also note a concentration-dependent effect of marimastat on target identification, where the higher concentration provided both a higher number of targets and less noise compared to the lower concentration of marimastat.

In addition to providing information about direct target interactions, PISA also allows for off-target effects to be investigated. Off-target binding of a drug may result in adverse effects that decrease (or complicate) its therapeutic utility (109,111), and small molecule drugs in particular tend to bind a myriad of molecular targets (112). For example, inhibitors of serine proteases exist that may be effective against medically significant snake venom serine proteases (SVSPs), but they may also cross-react with endogenous serine proteases in human plasma, which are critical for normal coagulation cascade activation (58). Our PISA analyses identified evidence of the interaction of marimastat with off-target toxin families, including CTLs and PLA_2_ toxins, which were also supported by our liquid chromatography and gel electrophoresis results. These findings are also consistent with prior studies that have shown marimastat and another SVMP inhibitor, prinomastat, can reduce PLA_2_-based anticoagulant venom effects (54). While CTLs are a minor and less clinically relevant component of *C. atrox* venom (95), PLA_2_s are more likely to be medically significant and tend to be fairly ubiquitous and abundant across diverse snake venoms (24,113–115). Though we did not detect any reduction in PLA_2_ activity in marimastat-treated samples (data not shown), off-target effects should be considered when investigating small molecule inhibitors of snake venom toxins, as they may demonstrate effects on other medically significant targets and/or contribute to unexpected outcomes *in vivo*.

Multiple snake venom gene families have undergone substantial gene family expansion, diversification, and neofunctionalization that has in many cases resulted in elevated rates of nonsynonymous substitutions in regions of these proteins that determine biological function (12). This trend has been observed in SVMPs (116), SVSPs (117), PLA_2_s (118,119), and 3FTXs (120), and has resulted in large multi-gene toxin families with similar structure but a wide array of biological functions and pharmacological effects which can also vary substantially across species (5,9,11,13,121). Indeed, this diversity of proteoforms within and across species presents an extreme challenge for the development of effective therapeutics to target the effects of these diverse and species-specific toxin cocktails. A major step to addressing this challenge has resulted in efforts to identify the most bioactive and medically relevant toxic proteins and proteoforms in venom using “omics” technologies, which has been referred to as “toxicovenomics” (122–125). A PISA-based approach in combination with toxicovenomics has the potential to take the key next step to address this complex problem through the screening of molecules that may neutralize the action of venom toxins across a wide variety of species that display high variability of medically significant venom toxin families, proteoforms, and activities. PISA and other high-throughput approaches provide promising paths forward for screening of large numbers of commonly studied and currently unexplored inhibitors against a wide scope of venoms for more rapid development of alternative snakebite therapies.

