## Supplementary figures and images for "ASSESSING TARGET SPECIFICITY OF THE SMALL MOLECULE INHIBITOR MARIMASTAT TO SNAKE VENOM TOXINS: A NOVEL APPLICATION OF THERMAL PROTEOME PROFILING"

### Supplemental Figure 1

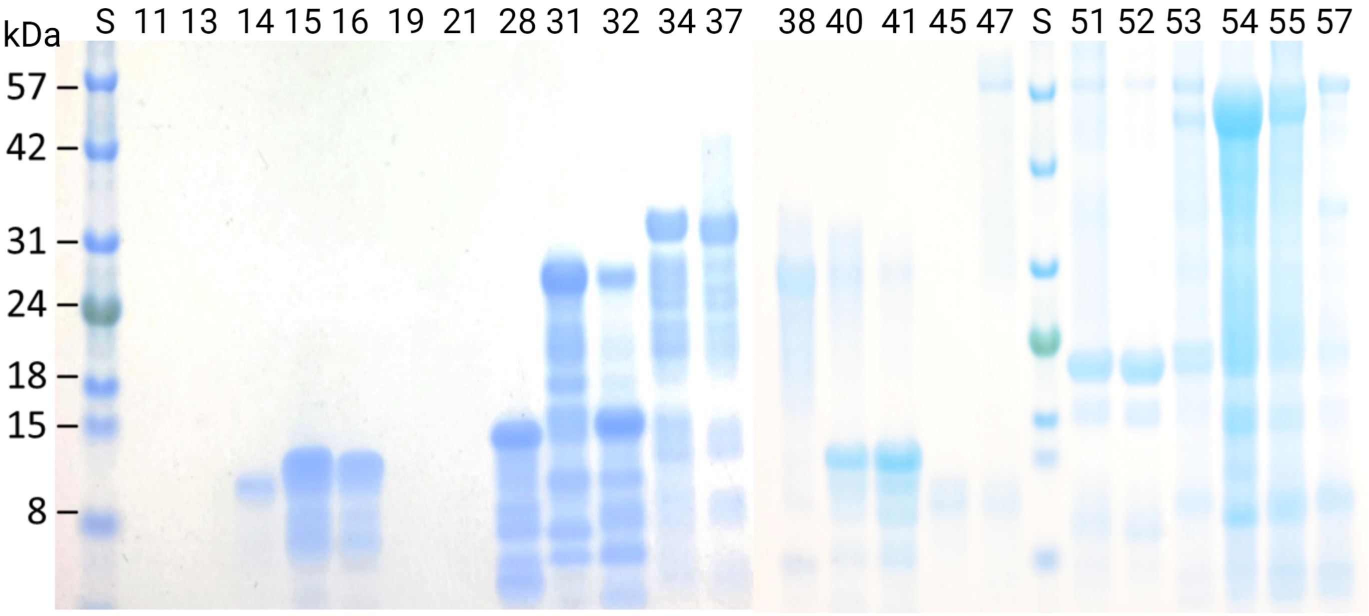
